# Shared neural mechanisms between imagined and perceived egocentric motion - A combined GVS and fMRI study

**DOI:** 10.1101/385625

**Authors:** Gianluca Macauda, Marius Moisa, Fred W. Mast, Christian C. Ruff, Lars Michels, Bigna Lenggenhager

## Abstract

Many cognitive and social processes involve mental simulations of a change in perspective. Behavioral studies suggest that such egocentric mental rotations rely on brain areas that are also involved in processing actual self-motion, thus depending on vestibular input. In a combined galvanic vestibular stimulation (GVS) and functional Magnetic Resonance Imaging (fMRI) study, we investigated the brain areas that underlie both simulated changes in self-location and the processing of vestibular stimulation within the same individuals. Participants performed an egocentric mental rotation task, an object-based mental rotation task, or a pure lateralization task during GVS or sham stimulation. At the neural level, we expected an overlap between brain areas activated during vestibular processing and egocentric mental rotation (against object-based mental rotation) within area OP2 and the Posterior Insular Cortex (PIC), two core brain regions involved in vestibular processing. The fMRI data showed a small overlap within area OP2 and a larger overlap within the PIC for both egocentric mental rotation against object-based mental rotation and vestibular processing. GVS did not influence the ability to perform egocentric mental rotation.

Our results provide evidence for shared neural mechanisms underlying perceived and simulated self-motion. We conclude that mental rotation of one’s body involves neural activity in the PIC and area OP2, but the behavioral results also suggest that those mental simulations of one’s body might be robust to modulatory input from vestibular stimulation.

## Introduction

Humans rely on mental simulations of action and perception to infer and evaluate the future state of one’s own body and its environment (Brecht, 2017; Hesslow, 2002; Moulton and Kosslyn, 2009). This is essential for the mental projection of the self to a different spatial location (egocentric mental rotation or mental perspective taking), which is also thought to be at the base of more complex social cognition (Kessler and Thomson, 2010; Wang et al., 2016; Zacks and Michelon, 2005).

Research in various sensory modalities has shown that the neural architecture for such mental simulations comprises similar neural pathways as those involved in actual perception and overt action (Kosslyn et al., 2001). In particular, it has been suggested that the vestibular system, which codes actual self-motion and self-orientation in space, is likely involved in mental changes of self-location (Ellis and Mast, 2017; Lenggenhager and Lopez, 2015; Mast et al., 2014). This is supported by several behavioral studies investigating the link between vestibular processing and egocentric mental rotation in patients with vestibular deficits (Candidi et al., 2013; Grabherr et al., 2011), but also in healthy participants during real or artificial vestibular stimulation (Deroualle et al., 2015; Dilda et al., 2011; Falconer and Mast, 2012; Ferrè et al., 2014; Grabherr et al., 2007; Grabherr and Mast, 2010; Lenggenhager et al., 2008; Pavlidou et al., 2017; van Elk and Blanke, 2014). Moreover, disturbances of perceived self-location are more common in vestibular patients (Lopez and Elzière, 2017) and often co-occur with vestibular illusions (Blanke, 2004; Blanke et al., 2004; Blanke and Mohr, 2005; Bonnier, 1893). Even out-of-body experiences (Blanke, 2004; Blanke et al., 2002) – possibly the most extreme form of illusory (and disembodied) change in egocentric perspective - have been repeatedly linked to vestibular processes (Blanke et al., 2004; Lopez and Elzière, 2017). However, while all this indirect evidence suggests shared neural mechanisms between vestibular processing and mental simulation of changes in self-location, no study until now has directly investigated this suggested shared neural overlap between egocentric mental rotation and vestibular processing.

Here, we used functional magnetic resonance imaging (fMRI) and galvanic vestibular stimulation (GVS) to locate brain areas that are involved in both vestibular processing and egocentric mental rotations. To this end, we adapted a mental rotation task (Keehner et al., 2006) that is similar to other tasks employed to study perspective taking (compare Kessler and Thomson, 2010). In this task, participants had to perform an egocentric mental rotation to a specified position to decide whether a certain object would be on their right or on their left. In the control condition, participants were presented the same stimuli but were instructed to perform an object-based mental rotation of the stimulus. Importantly, we hypothesized that egocentric mental rotations rely more on vestibular processing, since allocentric rotations, as necessary for the control condition, do not involve a mental change of one’s own position in space (Wang et al., 2016). To directly modulate activity of the individuals’ vestibular cortex and thus possibly influence participants’ ability to perform egocentric mental rotations, we used GVS. We hypothesized to find an overlap of activity induced by the processing of vestibular information elicited by GVS and egocentric mental rotation. In contrast to other modalities, there is no consensus whether a primary vestibular cortex exists, since vestibular signals are multisensory already at an early stage of processing (Lopez and Blanke, 2011). Nevertheless, two recent meta-analyses of neuroimaging studies investigating the neural correlates of artificial vestibular stimulation locate a core area of vestibular processing in the posterior insula and the parietal operculum, referred to as area OP2 (Lopez et al., 2012; zu Eulenburg et al., 2012). Area OP2 has been suggested to be the homologue to the parieto-insular vestibular cortex (PIVC) in non-human primates (Eickhoff et al., 2006). Yet, this definition of the PIVC has not been used consistently over different vestibular neuroimaging studies (Frank and Greenlee, 2018). In fact, there is evidence that the area immediately posterior to the PIVC receives vestibular input as well, but is also activated by visual motion cues such as optic flow (Billington and Smith, 2015; Frank et al., 2014, 2016b). This area has been labeled posterior insular cortex (PIC) and differs from the PIVC anatomically and functionally (Frank et al., 2014, 2016b), and in terms of functional (Smith et al., 2018) and anatomical (Wirth et al., 2018) connectivity. With a few recent exceptions, previous neuroimaging studies did not distinguish between PIC and PIVC (Frank and Greenlee, 2018; Lopez et al., 2012; zu Eulenburg et al., 2012).

We therefore selected area OP2 and PIC as our regions of interest for the analysis of the neural overlap between vestibular processing and egocentric mental rotation. Regarding the behavioral data, we expected that GVS interferes with egocentric more than allocentric mental rotation.

## Methods

### Participants

Thirty-one male, right-handed participants with no history of neurological, psychiatric or vestibular disorders took part in the first part of the study. All participants had normal or corrected-to-normal vision. From those, twenty healthy male participants (mean age = 25.55, SD = 5.4, range = 20-40 years) were selected for the fMRI task, based on the criterion that they perceived motion sensations during the sinusoidal GVS below a maximal amplitude of 2 mA. Moreover, participants needed to be able to solve the egocentric and object mental rotation task, and needed to be right-handed because there is evidence for dominance in cortical vestibular processing for the non-dominant hemisphere (Dieterich et al., 2003). All participants gave written informed consent before the study. The study was approved by the Cantonal ethics committee of Zurich. Participants were financially compensated for their participation in the behavioral and fMRI experiment.

### Task

The mental rotation task was adapted from Keehner and colleagues (2006). The presentation of the visual stimuli, the GVS and the response recording were programmed in Cogent (Wellcome Trust Centre for Neuroimaging, University College London, London, UK; http://www.vislab.ucl.ac.uk/cogent.php) implemented in Matlab (MathWorks). The visual stimuli were created in Blender, a free 3D creation suite (https://www.blender.org/) according to the parameters given in Keehner and colleagues (2006). They consisted of a circular table with a ball on top viewed from a 45° angle (see figure 1A). Moreover, an arrow below the table indicated the direction and distance of the mental rotation that had to be performed. The arrow length was set to 90°, 120° and 150° for left and right directions. Additionally, control trials were created where the arrow indicated no mental rotation (see also figure 1B). The ball position on the table varied for each arrow length in intervals of 30°. The task consisted of three different instructions. In the *egocentric mental rotation* condition, participants were instructed to imagine that the table remained stationary, while they mentally moved themselves around the table following the arrow to the tip, facing the center of the table from this new spatial location. In the *object mental rotation* condition, participants were instructed to imagine that their position remained stationary while they mentally rotated the table with the ball on it by the distance and in the direction indicated by the arrow. In the *no rotation* condition, participants had to perform no mental rotation. In all conditions, participants had to decide whether the ball was on their left or right after they performed the mental rotation (except for the *no rotation condition*) as fast and as accurately as possible. Responses were given with the right index finger (‘ball on left’) or right middle finger (‘ball on right’).

**Figure 1.**
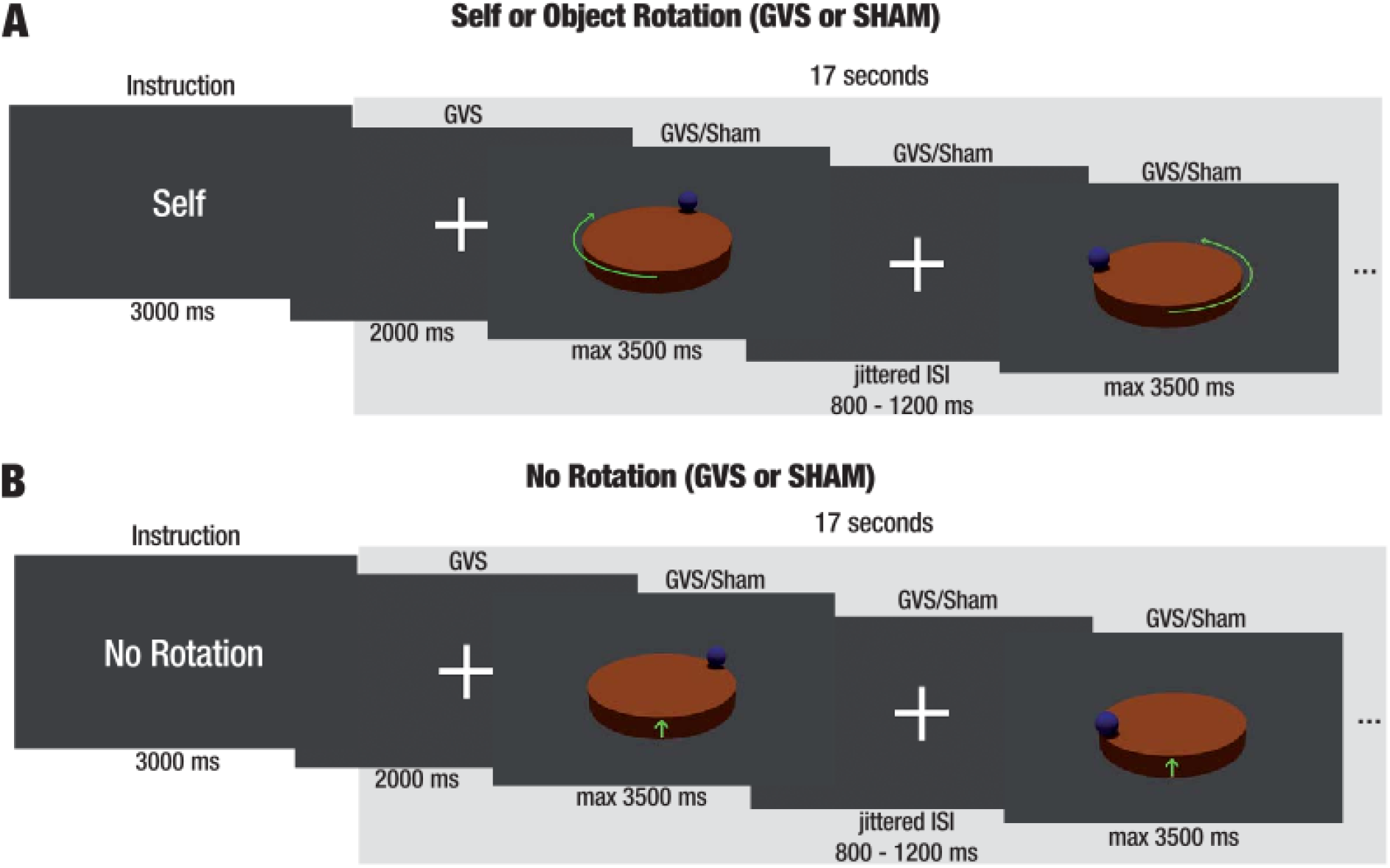
A depiction of the fMRI task for two example blocks. Figure **1A** illustrates the instruction, example trials and durations for an egocentric mental rotation block with either GVS or sham stimulation. Each block lasted 20 seconds. For every trial participants had to either mentally rotate their own position to the top of the arrow (egocentric condition) or to rotate the table with the ball on it in the direction of the arrow (object condition). They were instructed to indicate the ball’s position after the mental rotation as fast and accurately as possible. Figure **1B** depicts example trials for the no rotation control condition, during which participants were instructed to indicate as fast and accurately as possible whether the ball was on the left or on the right.

Importantly, visual stimuli and responses were identical for the egocentric and object mental rotation condition. Control trials differed only in the absence of the arrows (the no rotation conditions were indicated by a green pointer, see above). The no rotation control trials served to compare activity without spatial rotation with the conditions when participants had to perform an egocentric or allocentric mental rotation. Participants had to follow the instruction as the incorrect mental rotation strategy would have resulted in an error rate of 50%.

### Procedure and Experimental Design

The study took place on two different days. On day one, participants were informed about the study and gave their written informed consent. Afterwards sinusoidal GVS was applied in a supine position, at 1 Hz and different intensities (up to a maximum of 2 mA peak-to-peak) to find the intensity at which vestibular sensations were elicited (see Lenggenhager et al., 2008 for a similar procedure). If GVS was painful or participants did not perceive any vestibular sensation, they were not invited to the fMRI experiment. The established GVS intensities elicited postural instability in all participants that perceived vestibular sensations. Participants also completed the egocentric and object mental rotation task. Unlike the task used during fMRI, the behavioral task during the first session also included angles of 30 and 60 degrees. If accuracy for the egocentric or object mental rotation task was below 70 percent, participants were not invited to the fMRI experiment.

Twenty eligible candidates were selected for the fMRI experiment. Participants performed additional practice trials of the mental rotation task outside the scanner. For the fMRI, the mental rotation task consisted of a within-subject 3 (rotation tasks: egocentric, object, no rotation) × 2 (GVS, sham) design. These six conditions were presented in blocks of 20 seconds. Each block started with the rotation instruction presented for 3 seconds. After that, a fixation cross was presented for 2 seconds while the GVS started. In the sham blocks, the GVS stopped after these 2 seconds, while in the GVS conditions the signal continued for the remaining 15 seconds. During this interval, randomized rotation stimuli were presented (described in *Task*) for a maximum of 3.5 seconds each. After a response was given, the visual stimulus disappeared and a fixation cross was presented for a random duration of 0.8 to 1.2 seconds. Thus, the number of presented stimuli per block depended on the participants’ speed and participants were constantly engaged in performing mental rotations within a block. The end of the block was followed by the next instruction. The six different conditions were pseudorandomized within a run, with each condition being presented four times per run (each run lasted approximately 9 minutes). Participants performed four runs in total. Between runs, participants were given the opportunity to rest. Moreover, participants were asked after each run whether the individually determined GVS signal still elicited vestibular sensations. If no vestibular sensations were reported, then the signal’s amplitude was slightly increased in steps of 0.1 mA until vestibular sensations were perceived again.

### Vestibular Stimulation

Galvanic vestibular stimulation was delivered by a bipolar MR-compatible battery-driven current stimulator (NeuroConn DC-Stimulator PLUS) positioned outside the MR-scanner room. MR-compatible circular electrodes (diameter 3 cm) were attached to the participants’ mastoid processes and connected to the stimulator by means of two RF filter modules and MR-compatible cables. The electrodes were fixated using conductive paste and fixation bandages. The vestibular stimulation consisted of sinusoidal alternating current (AC) passed between the two electrodes at 1 Hz, in line with previous fMRI studies that used GVS (Smith et al., 2011). Sinusoidal GVS at 1 Hz has been shown to induce the strongest vestibular sensations (Stephan et al., 2005). The amplitudes were set individually according to the behavioral pretest session and the experienced vestibular sensations in the MR-scanner, but never exceeded 2 mA peak-to-peak. The sinusoidal stimulation elicited a sensation of sinusoidal roll motion of the head in the naso-occipital axis in all participants. The start of the GVS signal was precisely synchronized to the visual stimuli via in-house software and Cogent implemented in Matlab <u>(the same setup as the one used in Moisa et al. 2016; and in Bächinger et al. 2017)</u>.

### Imaging parameters

MR images were obtained using a Philips Achieva 3T whole-body MR scanner equipped with an eight-channel MR head coil. Each of the four experimental runs consisted of 216 volumes (voxel size, 2.5 × 2.5 × 3 mm^3^; 0.5 mm gap; matrix size, 96 × 96; repetition time (TR) 2610 ms; echo time (TE) 30 ms; 40 slices acquired in ascending order). T1-weighted multislice fast-field echo B0 scans were acquired for correction of possible static distortion produced by the presence of the GVS electrodes (voxel size, 3 × 3 × 3 mm^3^; 0.75 mm gap; TR/TE1/TE2 403/4.1/7.1 ms; flip angle, 44°; no parallel imaging; 37 slices). A high-resolution T1-weighted 3D fast-field echo structural scan was also acquired for image registration during post-processing (181 sagittal slices; matrix size, 256 256; voxel size, 1 mm^3^; TR/TE/inversion time (TI) 8.0/3.7/181 ms).

### Data Analysis

#### Behavioral

Behavioral data and regression parameters extracted from the fMRI models were analyzed with R (R Core Team, 2013). Bayesian multilevel models were calculated using the R-package brms (Bürkner, 2016) based on rstan (Guo et al., 2016). *Post-hoc* Bayesian correlations robust to outliers were calculated using a correlation model implemented in rstan (Bååth, 2013; Baez-Ortega, 2018). This correlation model is made robust to outliers by replacing an assumed bivariate normal distribution with a bivariate t-distribution.

Bayesian procedures provide posterior probability distributions for all estimated parameters. Non-informative priors were used for all parameters. Samples of each parameters’ posterior distribution were drawn with a Hamiltonian Monte Carlo sampling algorithm implemented in Stan (Carpenter et al., 2017). Samples were generated by four independent Markov chains, each with 1000 warm-up samples, followed by another 1000 samples drawn from the posterior distribution. For each Markov Chain, the last 1000 samples were saved for further statistical inference. To confirm that the samples for each chain converged to the same posterior distribution, R-Hat statistics were calculated (Gelman et al., 2014). For all calculated models, the R-Hat statistics were below 1.01, reflecting a low ratio of variance between the four chains to the variance within the chains. In addition, the visually inspected chains indicated that all Markov chains converged to the same posterior distribution of the estimated parameters. The 95% Bayesian credible intervals (CI) of these posterior distributions can be interpreted as the probable range of the parameter given the data and the model. The existence of an effect is inferred if the CI does not contain zero.

To test possible effects of GVS on the accuracy of mental rotation, we employed a Bayesian multilevel logistic regression with the correct responses as dependent variable. As we were interested in the interaction of the stimulation (GVS, sham) and the mental-rotation task (egocentric, object), the no-rotation control trials were not included in this model. The model consisted of four separate parameters on the population level for each condition defined by the experimental factors of stimulation (GVS vs. sham) and mental rotation task (egocentric vs. object). Moreover, the maximal random-effects structure justified by the experimental design was implemented(Barr et al., 2013), resulting in by-participant random effects for each condition.

To investigate effects of GVS on the response times in the egocentric mental rotation condition, a Bayesian multilevel multiple regression with a lognormal likelihood function was modelled. Similar to the accuracy model, we were interested in the interaction of the stimulation (GVS, sham) and the mental rotation task (egocentric, object) and thus did not model the data for the no-rotation control trials. As for the accuracy model, the reaction model consisted of four separate parameters on the population level for each condition defined by the experimental factors of stimulation (GVS vs. sham) and mental rotation task (egocentric vs. object), as well as by-participant random effects for each condition.

### fMRI - GLM

FMRI data were preprocessed and analyzed with Statistical Parametric Mapping (SPM12, http://www.fil.ion.ucl.ac.uk/spm) and additional toolboxes implemented in Matlab (MathWorks, Natick, Massachusetts, USA). Functional images were first corrected for geometric distortions using subject-specific field maps. Moreover, functional images were motion-corrected to the first image, slice-time corrected, normalized to MNI space using the segmentation parameters of the participants’ structural image, spatially resampled to 3 mm isotropic voxels and spatially smoothed with an isotropic Gaussian full-width at half-maximum kernel (FWHM) of 6 mm. A temporal high-pass filter (128 s cut off) was applied to remove low frequency drifts.

Statistical analysis was performed on two levels: In the first-level analysis, a subject-specific General Linear Model (GLM) was calculated. The first-level design matrix included fixed effects over all four runs. The model included six event-related main regressors for each run, one for each condition resulting from the 3 × 2 design. Importantly, only correct trials were included in the analysis. Reaction times were included as parametric regressors. An additional regressor modelled the onset of the vestibular stimulation at the beginning of each block. Regressors of interest were convolved with a canonical hemodynamic response function. Six participant-specific head motion parameters were included as regressors of no interest to control for BOLD signal changes that correlated with head movements. Contrast images were created for all regressors of interest.

To identify brain regions that were involved in egocentric mental rotation, the contrast *egocentric rotation sham > object rotation sham* was calculated for each participant. Moreover, to identify brain regions involved in vestibular processing independently of a mental rotation the contrast *no rotation during GVS > no rotation during sham* was calculated. To identify brain regions that were affected differently by the interaction of the mental rotation task and the stimulation, the interaction contrasts *(egocentric mental rotation GVS > egocentric mental rotation sham) > (object mental rotation GVS > object mental rotation sham) and (object mental rotation GVS > object mental rotation sham) > (egocentric mental rotation GVS > egocentric mental rotation sham)* were calculated. All these contrasts of interest were extracted for group level analysis.

All group analyses were calculated non-parametrically and performed in the Statistical Non-Parametric Mapping toolbox (SnPM version 13, http://warwick.ac.uk/snpm). SnPM uses a permutation approach and thus makes fewer assumptions about the underlying distribution of the data (Nichols and Holmes, 2002). In these second-level analyses, voxel-level pseudo-T statistics were obtained using variance smoothing with an isotropic Gaussian FWHM kernel of 6 mm and 5000 permutations. For the analyses within the OP2 and the PIC, these areas were included as explicit masks (see also Figure 2). Only voxel-level FWE corrected p-values below 0.05 are reported, except for the conjunction analysis at the whole brain level, where we also report the voxel-level uncorrected p-values below 0.001 to provide the reader with an overview of the whole brain results of the study’s main aim. However, due to the lack of correction for multiple comparisons, those results will not be interpreted. The anatomical labelling was performed with the help of the xjView toolbox (http://www.alivelearn.net/xjview) and the Anatomy toolbox (Eickhoff et al., 2005).

**Figure 2.**
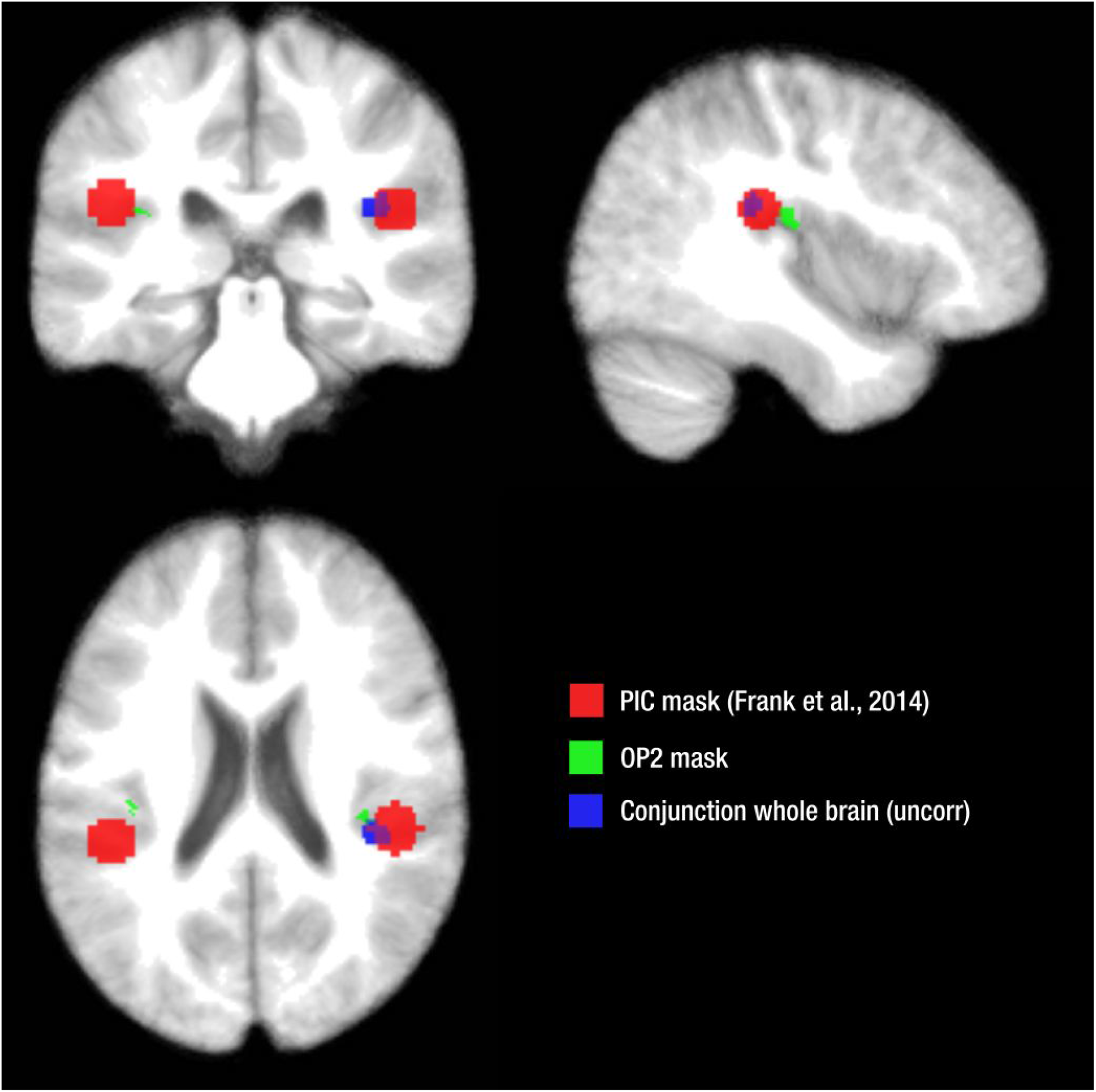
In red (PIC) and green (OP2) the masks used for the conjunction analysis. The PIC mask was created on the basis of the found by Frank et al. (2014). The OP2 mask was created with the Anatomy Toolbox. In blue, the whole brain conjunction analysis at *p < .001, uncorrected*. The figure was created in MRIcron (Rorden and Brett, 2000).

To identify voxels within the vestibular areas that are involved both in vestibular processing and in egocentric mental rotation, non-parametric conjunctions for the contrasts *no rotation GVS > no rotation sham* and *egocentric mental rotation sham > object mental rotation sham* were calculated on the second level on the whole-brain level and within *a priori* defined masks of the area OP2 and the PIC. While the mask of the area OP2 was created in the Anatomy Toolbox (Eickhoff et al., 2005), the mask of the PIC was based on coordinates of previous literature (Frank et al., 2014) and consisted of circular shapes of 8mm around the Talairach-coordinates *x = −44, y = −32, z = 22* for the left PIC and *x = 46, y = −29, z = 20* for the right PIC. These coordinates were transformed to MNI-space. For further exploratory analyses, individual mean parameter estimates for each condition were extracted from the significant voxels.

Non-parametric group level analyses were calculated to reveal brain areas involved in vestibular processing (contrast *no rotation GVS > no rotation sham*) and the interaction of mental rotation task and vestibular stimulation (contrasts *[egocentric mental rotation GVS > egocentric mental rotation sham] > [object mental rotation GVS > object mental rotation sham] and [object mental rotation GVS > object mental rotation sham] > [egocentric mental rotation GVS > egocentric mental rotation sham]*). We also calculated a group level analysis for the main effect of GVS (*GVS conditions > sham conditions*) over all rotation tasks (*egocentric, object and no rotation*) to test whether the activity pattern in the current study is comparable to other studies that used GVS in fMRI (Lopez et al., 2012 for meta-analyses; see zu Eulenburg et al., 2012).

### Psycho-Physiological Interaction

As the classical GLM analysis revealed GVS induced activity in the expected vestibular core areas, the right PIC and right OP2 (see Results section), we further explored altered functional coupling to these areas during egocentric rotation as compared to object rotation in the sham stimulation, as well as during GVS versus sham stimulation in the no rotation task. Functional connectivity was quantified by the means of psychophysiological interaction (PPI) analyses with the right PIC and right area OP2 as seed regions (Friston et al., 1997). In each participant and each run, the physiological times series in the individual seed regions were extracted from the right PIC and right area OP2. For the right area OP2 we selected the right hemisphere of the anatomically defined mask used for the GLM. For the right PIC we build a circular shape with a radius of 8mm around the coordinates in the right hemisphere reported above. Within this masks a 4mm sphere was created around the peak activity at *P < 1* for each run in each participant for the contrast *no rotation GVS > no rotation sham* to detect areas activated by vestibular processing. The individual seed regions were defined as the overlap of the anatomical map and the shape around the peak activity. The psychological regressors consisted of the six different types of conditions (the combination of the rotation strategies and stimulation), the reaction time as parametric modulators and the onset of the GVS stimulation (see description of GLM model). Moreover, six psychophysiological regressors were generated as the combination of the six different types of conditions and the timeseries of the right PIC and right area OP2. Based on these six psychophysiological regressors we calculated the two contrasts of interest (*egocentric rotation sham versus object rotation sham* and *no rotation GVS versus no rotation sham*). Head motion parameters from the realignment were again included as regressors of no interest. The contrasts of interest were extracted for non-parametric group-level analyses at the whole brain level.

### Behavioral Results

#### Accuracy and Response Times

Figure 3A shows the group level proportion of correct answers, the 95% CI, as well as the individual proportion of correct answers. Bayesian multilevel logistic regression revealed an effect of the mental rotation task (difference of mean parameter estimates [MPE] on logit scale =.66, 95% CI = [.15, 1.21]), indicating more correct responses in the egocentric mental rotation trials, no effect of GVS (difference of MPE on logit scale = −.31, 95% CI = [−.70,.05]), and no interaction between rotation task and stimulation (difference of MPE on logit scale = .07, 95% CI = [−.66,.81]) for the proportion of correct responses.

**Figure 3.**
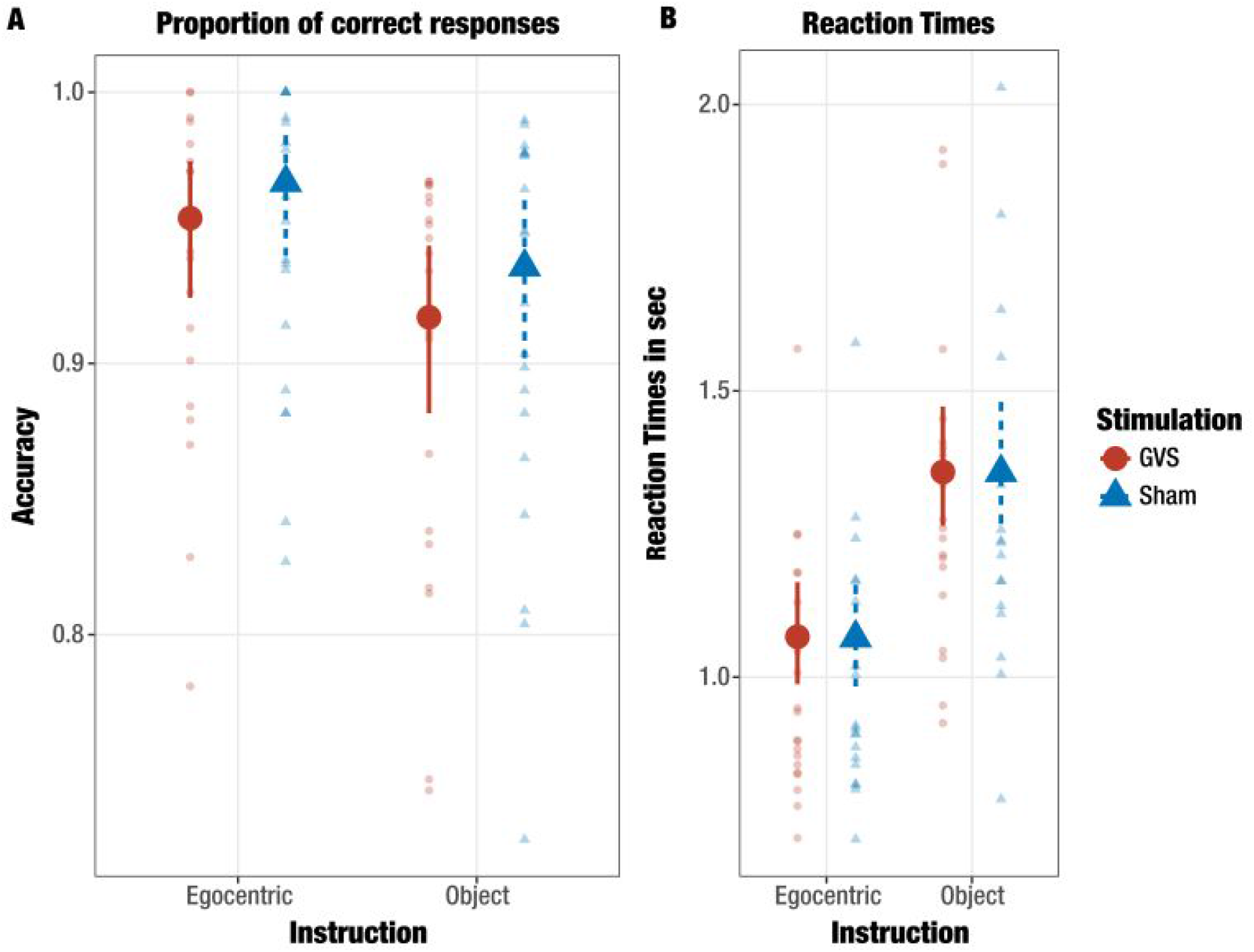
A summary of the behavioral results. In figure 3A, participants’ proportion of correct responses for the different conditions. The big dots show the group level parameter estimate of the GVS conditions from the multilevel logistic regression while the red lines indicate the 95% CI. The transparent smaller dots show the mean proportion of correct responses for each participant. Likewise, the triangles and the dashed lines indicate parameter estimates, CI, and individual proportion of correct responses for the sham conditions. Importantly, the analysis revealed a meaningful influence of the rotation task on the proportion of correct responses, but no influence of stimulation and no interaction. In figure 3B, the results of the reaction times analysis. As for the logistic regression, the big dots and blue triangles show the transformed parameter estimates of the mean from the multilevel lognormal regression analysis for the reaction times. The solid lines and the dashed lines indicate the 95% CI for the GVS and sham conditions, respectively. The small transparent dots and triangles represent individual median reaction times for each condition. Importantly, only correct responses were included in the analysis. The analysis revealed faster reaction times in the egocentric rotation trials, but no effect of GVS and no interaction.

Figure 3B shows the transformed parameter estimates from the Bayesian multilevel regression for the reaction times, the 95% CI, as well as the individual median reaction times for each condition. The Bayesian multilevel regression showed an effect of the mental rotation task (difference of MPE =.24, 95% CI = [.16, .33]), indicating that faster responses were given in the egocentric mental rotation condition, but no effect of GVS (difference of MPE = 0, 95% CI = [−.02, 02]) and no interaction between the rotation task and stimulation (difference of MPE = 0, 95% CI = [−.06, 05]).

### fMRI Results

#### Conjunction analysis

To test the hypothesis that vestibular processing and egocentric mental rotation both rely on shared areas within the vestibular cortex, a conjunction analysis on the group level was calculated for the first level contrasts *egocentric mental rotation sham > object rotation sham* and *no rotation GVS > no rotation sham*. The analysis at the whole brain level with a threshold of *p < .001*, not corrected for multiple comparisons revealed an activation cluster in the right Rolandic operculum and the right middle cingulum (see also Table 1 and Figure 2). The same conjunction analysis was also calculated within predefined masks of the area OP2 and the PIC. These analyses revealed a significant overlap in the right area OP2 (FWE voxel level threshold small volume corrected (svc), *peak p_FWE_* = 0.005, *k* = 3, *pseudo-t-peak* = 4.89, peak MNI coordinates 38, −28, 20) and bilateral PIC (FWE peak threshold small volume corrected (svc), *peak p_FWE_* = 0.003, *k* = 39, *pseudo-t-peak* = 4.33, peak MNI coordinates 42, −32, 20), see also figure 4A and B.

**Figure 4.**
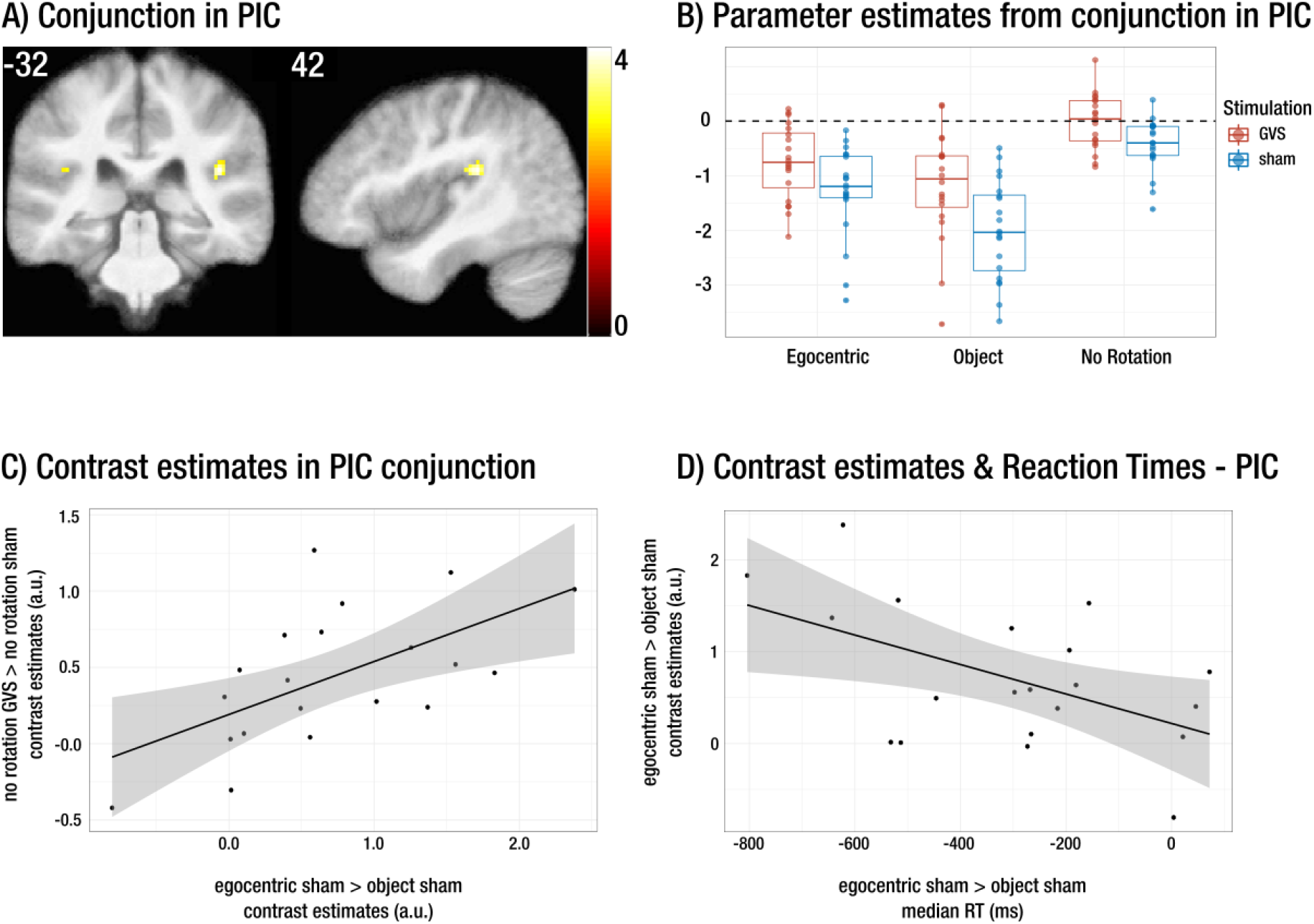
**A)** The results from the conjunction analysis for the contrasts *egocentric sham > object sham* & *no rotation GVS > no rotation sham* within the area PIC presented at a peak level threshold of p < 0.05 FWE-SVC corrected on the average T1-weighted image over all participants. The color scale indicates the Pseudo T-values. **B)** Boxplots for the mean parameter estimates for the cluster presented on the left for each participant. GVS conditions are presented in dark grey, sham conditions in lighter grey. Next to the boxplots, the dots represent the individual mean parameter estimates. This figure shows that the activation within the PIC cluster is higher for the egocentric mental rotation than for the object mental rotation condition. Moreover, there is also significantly higher activation for no rotation GVS compared to no rotation sham stimulation. **C)** The explorative *post-hoc* correlation analysis of mean parameter estimates extracted from the cluster presented in the panel A. The positive correlation indicates that participants with more activation in the egocentric sham condition also show more activation in the no rotation GVS condition. **D)** A *post-hoc* correlation analysis between the mean contrast estimates for the contrast *egocentric sham > object sham*, extracted from the cluster presented in panel A, and the correct median reaction times difference for the contrast *egocentric sham > object sham*. The negative correlation indicates that the faster the participants responded in the egocentric as compared to the object condition, the higher the positive activation difference within the cluster in the PIC.

**Figure 5.**
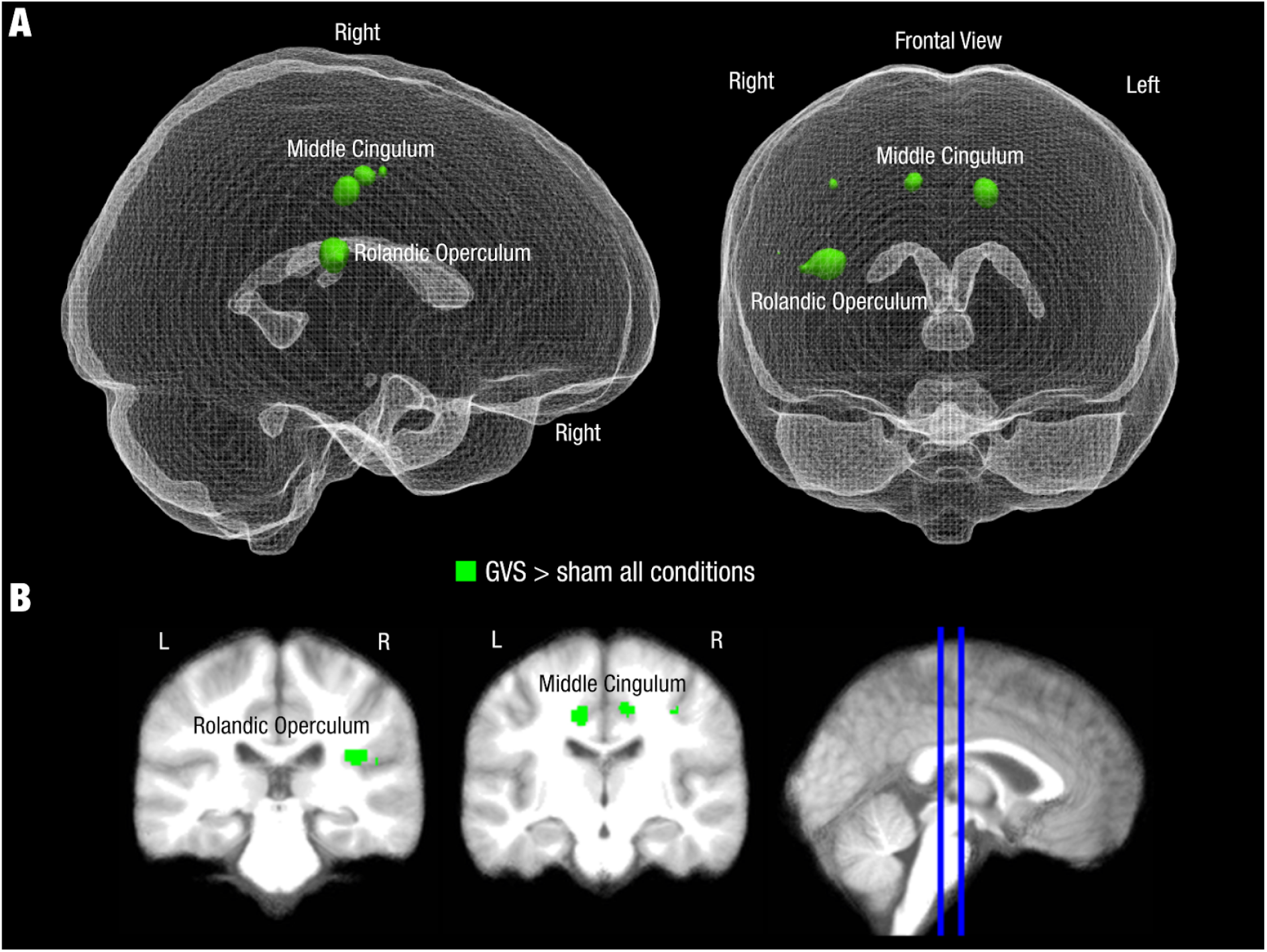
**A)** A rendering of the voxel-level FWE corrected non-parametric activations for the contrast GVS > sham over all rotation tasks presented on a glass brain (Madan, 2015). The upper left 3D rendering is the side view from the right side and the right rendering shows a frontal view. **B)** Coronal slices of the networks are illustrated to show the same activations. The coronal slices were created in MRIcron (Rorden and Brett, 2000). For more details see table 1.

**Table 1.** The results from the non-parametric group level fMRI analysis

| Region | X | Y | Z | Number of voxels | Pseudo T | Peak p |
| --- | --- | --- | --- | --- | --- | --- |
| **Conjunction egocentric rotation sham > object rotation sham, whole brain** |  |  |  |  |  |  |
| Right middle cingulate | 14 | -20 | 46 | 11 | 3.58 | > .001, uncorr. |
| Right Rolandic Operculum | 40 | -32 | 18 | 37 | 4.50 | > .001, uncorr. |
| **Conjunction egocentric rotation sham > object rotation sham within PIC** |  |  |  |  |  |  |
| Right PIC | 42 | -32 | 20 | 39 | 4.39 | .003, SVC-FWE |
| Left PIC | -38 | -32 | 20 | 2 | 3.23 | .034, SVC-FWE |
| **Conjunction egocentric rotation sham > object rotation sham within OP2** |  |  |  |  |  |  |
| Right OP2 | 38 | -28 | 20 | 3 | 4.89 | .005, SVC-FWE |
| **no rotation GVS > no rotation sham, whole brain** |  |  |  |  |  |  |
| Right Rolandic Operculum | 38 | -30 | 20 | 6 | 5.57 | .019, SVC-FWE |
| **GVS > sham overall, whole brain** |  |  |  |  |  |  |
| Right Rolandic Operculum | 38 | -32 | 18 | 31 | 7.97 | > .001, SVC-FWE |
| Left Middle Cingulum | -12 | -18 | 40 | 17 | 7.63 | > .001, SVC-FWE |
| Left Middle Cingulum | 12 | -18 | 44 | 122 | 6.56 | > .001, SVC-FWE |
| Right Precentral Gyrus | 36 | -16 | 44 | 54 | 5.72 | > .001, SVC-FWE |
| Right Cuneus | 16 | -76 | 32 | 12 | 5.48 | > .001, SVC-FWE |
| Left Insula | -34 | -32 | 22 | 46 | 5.10 | > .001, SVC-FWE |
| **GVS > sham overall within PIC** |  |  |  |  |  |  |
| Right PIC | 38 | -30 | 20 | 63 | 6.90 | > 0.001, SVC-FWE |
| Left PIC | -38 | -32 | 20 | 229 | 4.92 | > 0.001, SVC-FWE |
| **GVS > sham overall within OP2** |  |  |  |  |  |  |
| Right OP2 | 36 | -28 | 20 | 16 | 6.91 | > 0.001, SVC-FWE |
| Left OP2 | -36 | -30 | 20 | 3 | 3.42 | 0.021, SVC-FWE |

*Post-hoc* analyses of the extracted mean contrast estimates of the clusters within the OP2 ROI and PIC ROI revealed no correlation for the OP2 cluster (*rho* = 0.14, 95% CI = [−.30,.57]) and a meaningful correlation for the PIC cluster (*rho* = 0.53, 95% CI = [.20,.83]) between the contrast estimates for the contrast *egocentric mental rotation sham > object rotation sham* and the contrast *no rotation GVS > no rotation sham*. This indicates that participants with more PIC activity during the egocentric compared to the object rotation task also showed more PIC activity during GVS compared to sham stimulation. However, this relationship was not found for the OP2 cluster.

To explore whether the difference in OP2 and PIC activity between the egocentric and object rotation during sham would be reflected at a behavioral level (in the difference of the median reaction times between these two conditions), we calculated a correlation between each participant’s contrast estimates extracted from OP2 and the PIC for the contrast *egocentric mental rotation sham > object rotation sham* and the difference of the participant’s median reaction times for correct egocentric versus correct object rotations in the sham condition. This exploratory analysis revealed a negative correlation for the cluster within OP2 (*rho* = −.46, CI = [−.79,−.07]) and the cluster within the PIC (*rho* = −.45, CI = [−.79,−.07]), indicating that participants who were faster during egocentric mental rotation compared to the object rotation condition also showed more relative activity for the egocentric mental rotation condition within the cluster located in OP2 and the PIC (figure 4D).

#### Interaction of rotation task and stimulation

The non-parametric group level analysis for the interaction contrasts *(egocentric mental rotation GVS > egocentric mental rotation sham) > (object mental rotation GVS > object mental rotation sham)* and *(object mental rotation GVS > object mental rotation sham) > (egocentric mental rotation GVS > egocentric mental rotation sham)* revealed no significant peak or cluster.

#### Galvanic Vestibular Stimulation

To identify brain areas activated by GVS, the contrast *no rotation GVS > no rotation sham* was calculated on the group level. This group level contrast revealed a cluster with a significant peak in the right parietal operculum (FWE-corrected peak threshold, *peak p_FWE_* = .019, *k* = 6, see also table 1). We did not calculate the main effect of GVS over all rotation tasks, as our hypothesis was that the rotation task could possibly interfere with the vestibular stimulation. However, *post-hoc* inspection of this main effect of GVS (*GVS conditions > sham conditions*) over all rotation tasks (egocentric, object and no rotation) showed significant activation differences *p < 0.05* at the voxel level FWE corrected on the whole brain level in the right rolandic operculum, the bilateral middle cingulate cortex, the right precentral gyrus, the right cuneus and the left insula, consistent with the results of previous studies (see also table 1 and figure 4). To explore this activity differences and to exclude false negative activation, the same and the inverse contrast were also calculated within the OP2 and PIC mask separately. The results of these analyses are presented in Table 1. Importantly, the contrast *GVS conditions > sham conditions* revealed significant activity differences within both, the area OP2 and the PIC, while the contrast *sham conditions > GVS conditions* did not reveal any significant voxel within area OP2 or the PIC.

#### PPI analyses

The PPI analyses for the selected seed regions (right PIC and right OP2) and the contrasts of interest (*egocentric sham versus object sham* & *no rotation GVS versus no rotation sham)* did not reveal any functional coupling that survived correction for multiple comparisons.

## Discussion

The idea that vestibular areas not only process physical motion of one’s body but also supply a computational mechanism for imagined changes of self-location has fueled research on vestibular cognition in the last couple of years (Ellis and Mast, 2017; Mast et al., 2014). Several studies have attempted to influence participants’ egocentric mental rotation ability through vestibular stimulation, with varying outcomes (Deroualle et al., 2015; Dilda et al., 2011; Falconer and Mast, 2012; Lenggenhager et al., 2008; van Elk and Blanke, 2014). Moreover, separate evidence from neuroimaging studies on vestibular processing and on egocentric mental rotation suggest an overlap in the underlying neural processes. However, actual evidence for this idea within the same participants, or even within the same study, has been lacking. In the current investigation, we identify the hypothesized neural overlaps between the processing of perceived self-motion induced by GVS and simulated self-motion.

### Overlapping brain areas in the vestibular cortex

The neuroimaging data indicate that both, egocentric mental rotation and vestibular processing recruit brain areas within the vestibular cortex, in the current study operationalized as area OP2 and the PIC following the results of previous vestibular neuroimaging meta-analyses (Lopez et al., 2012; zu Eulenburg et al., 2012) and new insights into the functional and anatomical complexity of the vestibular cortex (Frank and Greenlee, 2018). In addition, the positive correlation between the PIC-BOLD contrast estimates for the effects of egocentric rotation versus object rotation and GVS suggests that the shared neural processes in this area are recruited to a similar degree by egocentric rotation and galvanic input. No correlation was found for the OP2 contrast estimates.

Indeed, the current investigation is the direct demonstration that vestibular brain areas are recruited more during egocentric mental rotation than during object rotation, in line with the large body of literature suggesting such an involvement (Candidi et al., 2013; Deroualle et al., 2015; Falconer and Mast, 2012; Grabherr et al., 2011; Grabherr and Mast, 2010; Lenggenhager et al., 2008; Mast et al., 2014; van Elk and Blanke, 2014). Only one previous study compared neural activity pattern during vestibular imagery and vestibular stimulation (zu Eulenburg et al., 2013). In that study, participants first underwent yaw rotations on a rotation chair and were instructed to recall the sensation of the experienced rotations afterwards during fMRI. The activations were compared to neural responses of vestibular processing induced by GVS. While GVS led to activity in the bilateral parietal operculum, bilateral supramarginal gyrus, inferior parietal lobule and other areas, the vestibular recall activated a network of brain areas involved in spatial referencing, motor processing and attention, but no significant activations or deactivations within areas known for processing vestibular information such as area OP2 or the PIC. Yet, it is unclear how participants solved the vestibular recall task in fMRI, as they were not asked about the strategy they used for the recall. The authors conclude that the high difficulty of vestibular recall might have prevented an intentional access of vestibular core areas. Moreover, the participants were exposed to the rotations while they were upright, and then later, when lying in the scanner, they had to recall the sensation in the supine position. The change in posture with respect to gravity makes it hard to comply with task instructions. In contrast, the present study used an established task of egocentric mental rotation and object mental rotation (Keehner et al., 2006). Our results suggest that the area that is processing vestibular information and is more involved in egocentric mental rotation relies on the area OP2 and PIC. The PIC has been shown to be responsive to artificial vestibular stimulation (Billington and Smith, 2015; Frank et al., 2014, 2016b) but also to visual motion (see Frank and Greenlee, 2018 for a recent overview). Due to its proximity to the PIVC, activity in the PIC has sometimes been misattributed to the PIVC. Only in the last couple of years, there has been evidence that the PIC is anatomically (Wirth et al., 2018) and functionally (Billington and Smith, 2015; Frank et al., 2014, 2016a) different. In contrast, the PIVC is suggested to be activated by vestibular stimulation and inhibited by visual stimulation. Moreover, the PIC is connected to regions that are associated with the perception of self-motion such as the CSv (Smith et al., 2011, 2017; Wall and Smith, 2008). The observed activity in the PIC rather than the PIVC in the present study could be due to nature of the task. In fact, participants had to mentally rotate themselves along an arrow that was presented visually. Thus, it cannot be excluded that the egocentric mental rotation relied on visuo-vestibular strategies despite the absence of visual motion cues.

Egocentric mental rotation is suggested to be an important mechanism in social perspective taking, which is performed on a daily basis and therefore is a highly trained ability (Kessler and Thomson, 2010). Thus, if perspective taking does in fact rely on areas involved in vestibular processing, it is reasonable to assume that an egocentric mental rotation task can activate vestibular core areas in a more subtle way. Additional evidence that the ability to perform egocentric mental rotations relies on the recruitment of vestibular core areas stems from the *post-hoc* correlational analysis between the brain activity during egocentric mental rotation and the corresponding median reaction times. In the present study, participants with more activity in the overlapping cluster within area OP2 and the PIC for the contrast *egocentric sham > object sham* were also faster during the egocentric as compared to object-based rotation during sham stimulation. We suggest that this increased activity in area OP2 and the PIC could be a predictor for the ability to perform egocentric mental rotations, though no causal inference can be made from the current data.

### No influence of GVS on behavior

Previous studies showed that GVS can influence egocentric mental rotation (Dilda et al., 2011; Lenggenhager et al., 2008) and perspective taking (Ferrè et al., 2014; Pavlidou et al., 2017). Studies using other artificial and natural vestibular stimulation techniques were further able to influence egocentric mental rotation (Falconer and Mast, 2012; Grabherr et al., 2007; van Elk and Blanke, 2014), and it has been shown that patients with vestibular disorders display a deteriorated ability to perform egocentric mental rotations (Candidi et al., 2013; Grabherr et al., 2011). In contrast, the results of this study show no influence of GVS on reaction times for egocentric and object mental rotation, and only a small influence on accuracy. In this context it is important to note that the present data show a difference in task difficulty between both tasks, with egocentric rotation being considerably easier (reaction times were faster and there was a higher proportion of correct responses). Similar results were obtained by Keehner and colleagues (2006) who used the same task. There is also no interaction of the rotation task and vestibular stimulation on the behavior level. Not surprisingly, a whole-brain analysis for the interaction of the rotation task and stimulation did not reveal any significant effects. One possibility to account for the absence of a behavioral effect is that the stimulation profile used in this study may not have been strong enough to interfere with egocentric mental rotation, especially in the supine position inherent to fMRI experiments. Given the recent advances in the flexibility and spatial precision of magnetoencephalography (Boto et al., 2018), future studies using this technique may potentially overcome this problem.

In addition, as egocentric mental rotation is an essential ability in daily life and vestibular processing is constantly ongoing, it seems reasonable to suggest that even though both these processes rely on shared mechanisms, there may be enough resources for processing both of them in parallel. The proportion of correct responses was very high, indicating low task difficulty. Because previous literature suggests a mental self-rotation strategy at angles above 90° (Kozhevnikov and Hegarty 2001), we only used three different angles of rotation. This may have reduced task difficulty and may have reduced a potential interference from concurrent GVS. However, the proportion of correct responses in the current study was comparable to the data presented by Keehner and colleagues (2006), who additionally used angles of 30° and 60°. Moreover, behavioral studies that found an effect of vestibular stimulation on egocentric mental rotation report similar proportions of correct responses, indicating similar levels of difficulty. It is important to note that the task was designed to allow disentangling an egocentric and allocentric mental rotation strategy with the identical visual stimuli. This was important for the analysis of the neuroimaging data, and - despite the absence of an influence of GVS - the behavioral data provide compelling evidence that participants used the two strategies as instructed.

### Vestibular activation

The present activation patterns elicited by GVS are comparable to other vestibular neuroimaging studies. Mainly, in the present study, GVS elicited activity in the right parietal operculum, more specifically in area OP2 and the Posterior Insula Cortex. This is in line with two recent meta-analyses (Lopez et al., 2012; zu Eulenburg et al., 2012) and a previous study that used GVS to delineate the primary human vestibular cortex and located it in the right hemisphere of area OP2 (Eickhoff et al., 2006). Our results also underline the predominance of the non-dominant right hemisphere for vestibular processing (e.g. Dieterich et al., 2003). The contrast of GVS versus sham over all rotation tasks revealed activity in the bilateral middle cingulate sulcus. Interestingly, similar activations in the bilateral cingulate sulcus elicited by GVS have been reported previously (Smith et al., 2011). This area has been detected by the same group as a response to egomotion induced by optic flow (Wall and Smith, 2008) and accordingly been labelled CSv. It is hypothesized that the CSv is involved in processing heading information (Wall and Smith, 2008) and potentially involved in visuo-vestibular interactions (Smith et al., 2017). Moreover, intracranial stimulation around this area has been shown to elicit vestibular sensations (Caruana et al., 2018). Interestingly, there is compelling evidence that there is anatomical and functional connectivity between the CSv and the PIC (Smith et al., 2018).

### Conclusion

Our results confirm recent ideas that vestibular brain areas are involved in egocentric mental rotation. The current study provides first evidence that both vestibular processing and egocentric mental rotation rely on overlapping activation within the vestibular cortex, specifically the PIC and area OP2. Part of the vestibular cortex is thus associated with the processes underlying mental self-rotation, demonstrating that vestibular areas are also involved when self-rotation is imagined while there is no vestibular sensory input causing the activation.

## Funding

This work was supported by the Swiss National Science Foundation (SNSF, grant numbers 142601 and 162480).

## Conflicts of interest

All authors declare that they have no conflict of interest.

